# Prospects for enhancing leaf photosynthetic capacity by manipulating mesophyll cell morphology

**DOI:** 10.1101/379065

**Authors:** Tao Ren, Sarathi M Weraduwage, Thomas D. Sharkey

## Abstract

Leaves are beautifully specialized organs designed to maximize the use of light and CO_2_ for photosynthesis. Engineering leaf anatomy therefore brings great potential to enhance photosynthetic capacity. Here we review the effect of the dominant leaf anatomical traits on leaf photosynthesis and confirm that a high chloroplast surface area exposed to intercellular airspace per unit leaf area (*S_c_*) is critical for efficient photosynthesis. The possibility of improving *S_c_* through appropriately increasing mesophyll cell density is further analyzed. The potential influences of modifying mesophyll cell morphology on CO_2_ diffusion, light distribution within the leaf, and other physiological processes are also discussed. Some potential target genes regulating leaf mesophyll cell proliferation and expansion are explored. Indeed, more comprehensive research is needed to understand how manipulating mesophyll cell morphology through editing the potential target genes impact leaf photosynthetic capacity and related physiological processes. This will pinpoint the targets for engineering leaf anatomy to maximize photosynthetic capacity.

**Highlight:** Cell morphology in leaves affects photosynthesis by controlling CO_2_ diffusion and light distribution. Recent work has uncovered genes that control cell size, shape, and number paving the way improved photosynthesis.

## Introduction

Increasing the conversion efficiency of intercepted radiation into biomass through enhancing photosynthesis has emerged as a promising focus to enhance crop production for meeting needs of the increasing population and for plant resilience in the face of the changing global climate (Long *et al*., 2015b; Ort *et al*., 2015; Souza *et al*., 2017). Some current approaches to improving photosynthesis in C3 crops, include conversion to C_4_ metabolism, engineering a photorespiratory bypass, optimizing regeneration of RuBP etc. It is estimated that these could increase the light conversion efficiency by 5%-60% compared with today’s best cultivars (Long *et al*., 2015b). Using the rice GRCROS model, Yin and Struik (2017) simulated nine genetic engineering strategies for enhancing C3 leaf photosynthesis to improve crop production under the present climate and the climate predicted for 2050. Although the knowledge of photosynthesis has exploded recently, there is no doubt that it is still a major challenge to understand the mechanisms, vulnerabilities, and potentials for improvement of photosynthesis (Niinemets *et al*., 2017).

Leaves are beautifully specialized organs evolved for efficient photosynthesis. Their morphological differences can explain substantial variations in photosynthetic properties among different species or the same species under different environmental constraints (Oguchi *et al*., 2005; Terashima *et al*., 2011; Peguero-Pina *et al*., 2017; Tsukaya, 2018). For example, alterations in photosynthetic performance of sun and shade leaves of the same plant are intimately connected with leaf anatomy (Terashima *et al*., 2001, 2006; Oguchi *et al*., 2005; Théroux-Rancourt and Gilbert, 2017). The inseparable relationship between leaf architecture and photosynthesis thus brings great potential to enhance photosynthetic efficiency by modifying leaf anatomy. Tholen *et al*. (2012) described how altering leaf anatomy could significantly influence light distribution and CO_2_ diffusion in the leaf. Nonetheless, precise targets for engineering leaf anatomy are still unknown because of poor understanding of the molecular mechanisms controlling leaf anatomy.

Leaf mass per unit leaf area (LMA), one important leaf morphological trait that affects photosynthesis, is also related to environmental stress and longevity of plants (Niinemets, 2001; Poorter *et al*., 2009; de la Riva *et al*., 2016). There exists considerable variation regarding the correlation between LMA and photosynthesis,. Negative relationships between LMA and photosynthesis are associated with higher investment in nonphotosynthetic tissue, such as cell walls (Onoda *et al*., 2004, 2017; Hassiotou *et al*., 2010) and lower mesophyll conductance (Flexas *et al*., 2008; Niinemets *et al*., 2009; Weraduwage *et al*., 2016). Positive relationships are associated with improvement of chloroplast exposure to intercellular airspace (Peguero Pina *et al*., 2017) and chloroplast CO_2_ concentration, which ultimately enhances the carboxylation rate (Cai *et al*., 2014; Flexas *et al*., 2014) under higher LMA owing to an increase in leaf thickness. The influence of LMA on photosynthesis depends in part on how LMA impacts mesophyll conductance. An increase in LMA related to increased leaf density might cause mesophyll cells to be densely packed, which will inevitably increase mesophyll CO_2_ resistance (Flexas *et al*., 2008; Niinemets *et al*., 2009; Weraduwage *et al*., 2016). However, if the mesophyll cell space is large enough to avoid excessive cell-to cell contact, it is possible to increase mesophyll and chloroplast surface area exposed to intercellular airspace per unit leaf area and improve mesophyll conductance (Peguero-Pina *et al*., 2017). Recently, John *et al*. (2017) wrote that LMA is determined by leaf cell numbers, dimensions and mass densities. Optimizing LMA through regulating mesophyll cell morphological features thus might be a good target to enhance leaf photosynthetic capacity.

Early in the 1960s and 1970s there were many attempts to correlate leaf anatomical characteristics with photosynthesis. Mesophyll cell size was found to be an important factor (Jellings and Leech, 1984). Wilson and Cooper (1970) reported that selection for a smaller cell size could be used to improve photosynthesis and yield in ryegrass (*Lolium perenne*). But at that time, it was not entirely known how mesophyll cell size impacts leaf photosynthesis rate; possible factors include cell volume, cell surface area, or the effect of cell shape on cellular packing within the leaf (Jellings and Leech, 1984). In the last two-decades, extensive investigations of leaf anatomical traits and photosynthesis among different species or same species under various environmental conditions have reemerged (Kogami *et al*., 2001; Hanba *et al*., 2002; Oguchi *et al*., 2005; Peguero-Pina *et al*., 2012, 2016; Sáez *et al*., 2017). Although the typical characteristics of leaves with high photosynthetic efficiency among different species or environmental factors are not always consistent, a higher proportion of chloroplast surface area exposed to intercellular airspace per unit leaf area (*S*_c_) is generally associated with high rates of photosynthesis. High *S*_c_ reduces the CO_2_ diffusion resistance in the leaf and improves CO_2_ concentration at the sites of carboxylation. Higher *S*_c_ values associated with more and smaller mesophyll cells could lead to selection for smaller cells. Achieving higher *S*_c_ values through manipulating mesophyll cell morphological traits might be a crucial spotlight of engineering leaf anatomy to enhance leaf photosynthetic efficiency.

Furthermore, increasing explorations of the key target genes regulating cell proliferation and expansion provides more possibilities for engineering leaf architecture to improve photosynthetic efficiency. Takai *et al*. (2013) revealed the gene *GREEN FOR PHOTOSYNTHESIS* (*GPS*) increases mesophyll cell number between vascular bundles, which leads to enhanced photosynthesis rates. Lehmeier *et al*. (2017) manipulated mesophyll cell density through promoting or repressing *KIP-RELATED PROTEIN1* (*KRP1*) and *RETINOBLASTOMA RELATED PROTEIN 1* (*RBR1*) gene to increase photosynthetic capacity. Our recent work also found that the leaves of *ROOT HAIR DEFECTIVE 3* (*RHD3*) defective mutants have more and smaller mesophyll cells, and their photosynthetic rate was much higher than wild type. All these indicate that regulation of mesophyll cell division, expansion and arrangement in leaves is a potential target of engineering leaf anatomy by genetic approaches to enhance photosynthetic efficiency. Therefore, this manuscript reviews the effects of the dominant leaf anatomical traits on leaf photosynthesis, analyzes the possibility of improving *S*_c_ through re-arranging mesophyll cells, and further discusses the potential influences of modifying leaf anatomy on CO_2_ diffusion, light utilization and other physiological processes. Finally, some potential target genes are explored.

## 2. Enhancing leaf photosynthetic capacity by altering leaf anatomy

### 2.1 Effects of leaf anatomical characteristics on photosynthesis

A typical C_3_ leaf cross-section, comprising two epidermal layers surrounding a tight array of columnar cells (palisades) and irregularly shaped cells (spongy), provides powerful tool to explore various morphological features associated with leaf physiological functions. Here we summarize 43 published research papers and extracted the major leaf anatomical and photosynthetic parameters (Table S1), including leaf thickness (*T*_L_), mesophyll layer thickness (*T*_mes_), fraction of intercellular airspace (*f*_ias_), *LMA*, mesophyll cell wall thickness (*T*_w_), chloroplast length (Lchl), chloroplast thickness (*T*_chl_), mesophyll surface area exposed to intercellular airspace (*S*_mes_), *S*_c_, stomatal conductance (*g*_s_), mesophyll conductance (*g*_m_) and photosynthesis rate (*A*_n_) (Fig. 1). There are huge variations in leaf morphological characteristics of different species in different environments, and the coefficient of variance (c.v.) values vary from 22% to 79%. The smallest coefficient of variation is the length of the chloroplast. The average chloroplast length is 4.81 μm, varying from 1.58 μm to 6.86 μm. The coefficient of variation of LMA is highest, averaging 76 g m^−2^ and ranging from 11 g m^−2^ to 285 g m^−2^.

**Fig. 1.**
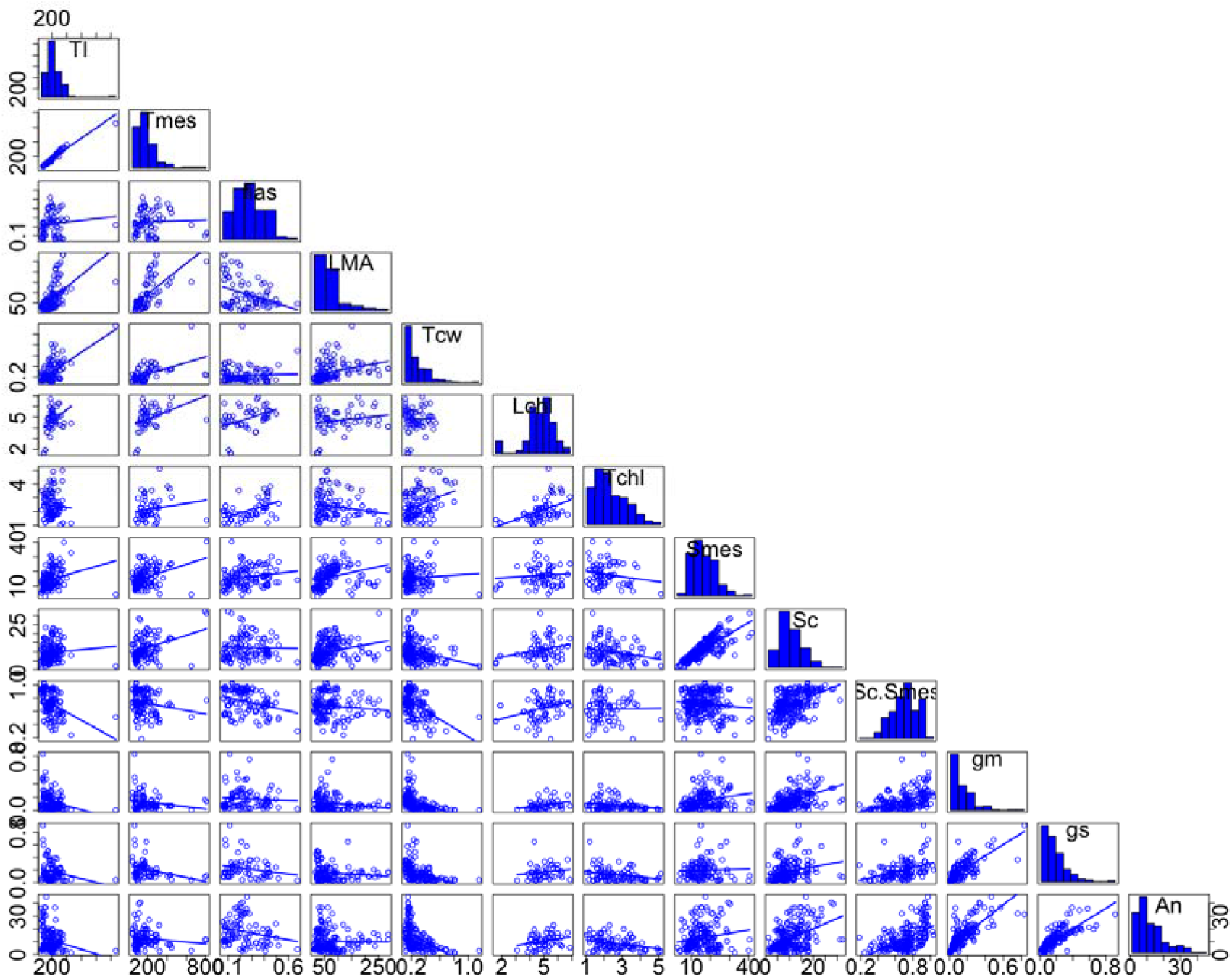
Scatterplot matrix of critical leaf morphology and photosynthetic parameters based on literature data. Note: The matrix was conducted by using the “car” package in R software. Diagonal panels show the distribution of each variable. Lower panels show scatterplot. *Tl*, leaf thickness; *Tmes*, mesophyll thickness; *fias*, fraction of intercellular air space; *LMA*, leaf dry mass per unit area; *Tcw*, mesophyll cell wall thickness; *Lchl*, chloroplast length; *Tchl*, chloroplast thickness; *Smes*, mesophyll surface area exposed to intercellular air space; *Sc*, chloroplast surface area exposed to intercellular air space; *Sc.Smes*, the ratio of Sc/Smes; *gm*, mesophyll conductance; *gs*, stomatal conductance; *An*, photosynthesis rate.

There is no significant correlation between *LMA* and *A*_n_ (Table 1), as reported in previous studies (Hassiotou *et al*., 2010; Veromann-Jürgenson *et al*., 2017). Moreover, the long distance between these two parameters indicates wide variability between *LMA* and *A*_n_ (Fig. 2). As decribed in many reports (Poorter *et al*., 2009; de la Riva *et al*., 2016; Veromann-Jürgenson *et al*., 2017; John *et al*., 2017), leaf thickness, mesophyll layer thickness, mesophyll cell wall thickness and intercellular airspace are the dominant factors influencing *LMA*. When the *LMA* value is less than 100 g m^−2^, an increase in *LMA* significantly promotes *S*_mes_ and *S*_c_ values. However, when it exceeds 100 g m^−2^, higher *LMA* value plays only a minor role in the improvement of the *S_mes_* and *S_c_* value. On the contrary, higher *LMA* reduces the *f_ias_* and *S_c_/S_mes_* values, possibly because of densely packed mesophyll cells, further decreasing *g_m_* (Table 2).

**Fig. 2.**
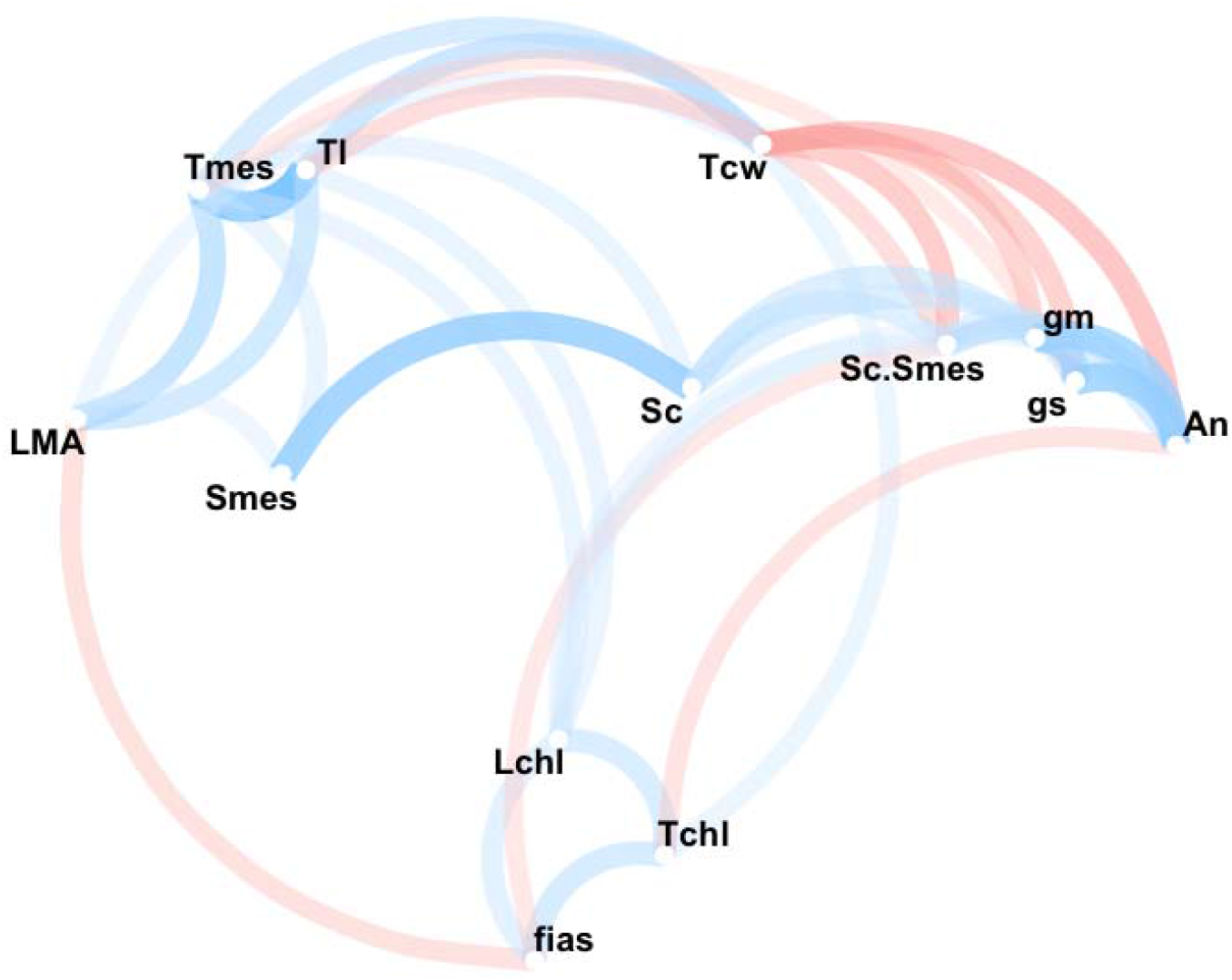
Correlation between leaf morphology and photosynthetic parameters based on literature data. Note: The correlation was conducted by using the “corrr” package in R software. The closer each variable is to another represents the higher relationship while the opposite is for widely spaced variables. The line color represents the direction of the correlation. The blue line is positive correlation and the red line is negative correlation. The line shade and thickness represent the strength of the relationship. The minimum correlation coefficient required to display a line between variables is 0.3.

**Table 1.** Matrix of correlation coefficient between leaf anatomical and photosynthetic parameters

|  | T<sub>L</sub> | T<sub>mes</sub> | f<sub>ias</sub> | LMA | T<sub>cw</sub> | L<sub>chl</sub> | T<sub>chl</sub> | S<sub>mes</sub> | S<sub>c</sub> | S<sub>c</sub>/S<sub>mes</sub> | g<sub>m</sub> | g<sub>s</sub> | A<sub>n</sub> |
| --- | --- | --- | --- | --- | --- | --- | --- | --- | --- | --- | --- | --- | --- |
| T<sub>L</sub> | 1.00 | 0.97*** | 0.11 | 0.55*** | 0.60*** | 0.36** | -0.062 | 0.24** | 0.085 | -0.38*** | -0.26** | -0.22* | -0.22** |
| T<sub>mes</sub> |  | 1.00 | 0.039 | 0.63*** | 0.52*** | 0.37** | 0.15 | 0.37*** | 0.36*** | -0.18* | -0.20* | -0.30** | -0.12 |
| f<sub>ias</sub> |  |  | 1.00 | -0.37*** | 0.033 | 0.42** | 0.52*** | 0.19* | -0.021 | -0.31*** | -0.065 | -0.22* | -0.23* |
| LMA |  |  |  | 1.00 | 0.33*** | 0.19 | -0.20* | 0.33*** | 0.27*** | -0.12 | -0.17* | -0.13 | -0.018 |
| T<sub>cw</sub> |  |  |  |  | 1.00 | 0.06 | 0.35*** | 0.071 | -0.27*** | -0.48*** | -0.44*** | -0.45*** | -0.52*** |
| L<sub>chl</sub> |  |  |  |  |  | 1.00 | 0.50*** | 0.081 | 0.27* | 0.32** | 0.32* | 0.16 | 0.26* |
| T<sub>chl</sub> |  |  |  |  |  |  | 1.00 | -0.18 | -0.26** | 0.017 | -0.24* | -0.28** | -0.34*** |
| S<sub>mes</sub> |  |  |  |  |  |  |  | 1.00 | 0.80*** | -0.099 | 0.22** | 0.03 | 0.18* |
| S<sub>c</sub> |  |  |  |  |  |  |  |  | 1.00 | 0.46*** | 0.49*** | 0.24** | 0.47*** |
| S<sub>c</sub>/S<sub>mes</sub> |  |  |  |  |  |  |  |  |  | 1.00 | 0.44*** | 0.41*** | 0.55*** |
| g<sub>m</sub> |  |  |  |  |  |  |  |  |  |  | 1.00 | 0.75*** | 0.76*** |
| g<sub>s</sub> |  |  |  |  |  |  |  |  |  |  |  | 1.00 | 0.78*** |
| A<sub>n</sub> |  |  |  |  |  |  |  |  |  |  |  |  | 1.00 |

Note: \*\*\*, \*\*, and \* represents significant at $P<0.001$ , $P<0.01$ and $P<0.05$ , respectively.

**Table 2.** Changes of leaf anatomical and photosynthetic parameters under different LMA

| LMA (g m<sup>-2</sup>) | <50 | 50-100 | >100 | P value |
| --- | --- | --- | --- | --- |
| T<sub>L</sub> (μm) | 145±61c | 187±67b | 320±172a | <0.001*** |
| T<sub>mes</sub> (μm) | 127±54b | 161±68b | 306±151a | <0.001*** |
| f<sub>ias</sub> | 0.276±0.165a | 0.298±0.108a | 0.186±0.128b | 0.0076** |
| T<sub>cw</sub> (μm) | 0.242±0.135c | 0.318±0.182b | 0.409±0.171a | <0.001*** |
| L<sub>chl</sub> (μm) | 4.38±1.53a | 4.79±0.60a | 4.91±0.88a | 0.26 |
| T<sub>chl</sub> (μm) | 2.38±0.88a | 2.60±0.95a | 2.09±0.78a | 0.081 |
| S<sub>mes</sub> (m<sup>2</sup> m<sup>-2</sup>) | 12.0±5.4b | 19.6±4.5a | 19.2±9.6a | <0.001*** |
| S<sub>c</sub> (m<sup>2</sup> m<sup>-2</sup>) | 7.9±4.1b | 12.6±5.1a | 11.3±5.7a | <0.001*** |
| S<sub>c</sub>/S<sub>mes</sub> | 0.68±0.15a | 0.72±0.18a | 0.61±0.16b | 0.013* |
| g<sub>m</sub> (mol m<sup>-2</sup> s<sup>-1</sup>) | 0.119±0.104a | 0.131±0.148a | 0.062±0.042b | 0.029* |
| g<sub>s</sub> (mol m<sup>-2</sup> s<sup>-1</sup>) | 0.177±0.147a | 0.123±0.080a | 0.134±0.118a | 0.073 |
| A<sub>n</sub> (μmol m<sup>-2</sup> s<sup>-1</sup>) | 10.9±8.3a | 9.6±7.6a | 9.8±5.5a | 0.54 |

Note: \*\*\*, \*\*, and \* represents significant at $P<0.001$ , $P<0.01$ and $P<0.05$ , respectively.

Despite different species and environmental conditions, significantly positive correlations between *S*_c_ and *A*_n_ and *g*_m_ are observed; *S*_c_ is the closest positive parameter to *A*_n_ and *g*_m_ of all summarized leaf morphological parameters. From our results, the average *S*_c_ value is 11.1 m^2^ m^−2^, ranging from 1.2 m^2^ m^−2^ of *Deschampsia antarctica Desv* in Lagotellerie island (Sáez *et al*., 2017) to 32.3 m^2^ m^−2^ of *Abies pinsapo* needles in the Mediterranean area (Peguero-Pina *et al*., 2012). High variations and significant positive correlation between *S*_c_ and *A*_n_ confirm that *S*_c_ is a potential target of engineering leaf anatomy to enhance leaf photosynthetic capacity.

### 2.2 Increase *S*_c_ by manipulating mesophyll cell morphology

*S*c is the most important leaf structural parameter in photosynthetic studies during the last decades. von Caemmerer and Evans (1991) showed that higher *S*_c_ could explain the relatively large CO_2_ transfer conductance of rice. Almost all subsequent studies have confirmed the importance of having a high *S*_c_ value for maximizing photosynthesis (Terashima *et al*., 2011). The *S*_c_ value is influenced by chloroplast number, distribution in the mesophyll and mesophyll surface area exposed to intercellular airspace. An increase in the number of chloroplasts in mesophyll cells is beneficial to improve Sc (Miyazawa and Terashima, 2001). Additionally, changes in light intensity and quality can induce movement and re-distribution of chloroplasts within cells, further affecting *S*_c_ (Hanba *et al*., 2002; Oguchi *et al*., 2003). The maximum value of *S*_c_ is determined by *S*_mes_, which is the closest positive parameter to *S*_c_ (Fig. 2). In our analysis the average *S*_c_/*S*_mes_ value is 0.71, implying that nearly 3/4 of mesophyll cell surface area facing intercellular airspace is occupied by chloroplast. For *Castanopsis sieboldii*, the *S*_c_/*S*_mes_ ratio could be 1.0 after 20 days of full leaf area expansion (Hanba *et al*., 2002). Oguchi *et al*. (2003) described that transferring the mature leaf from low to high light significantly increased photosynthesis rate, associated with improving *S*_c_. The photosynthesis rate in the low-light leaf was significantly lower than the leaf exposed to high light because of restricted *S*_mes_ value. Greater *S*_mes_ is a prerequisite for a larger *S*_c_. Therefore, simultaneously raising the *S*_mes_ and *S*_c_ is the key to enhance leaf photosynthetic efficiency.

Nobel *et al*. (1975) first reported in that the mesophyll cell surface area per unit leaf area (*A*_mes_/*A* in their notation), calculated assuming that mesophyll cells were cylindrical with hemispheres at each end, was correlated with CO_2_ assimilation rate. Evans *et al*. (1994) optimized the calculation depending on the shape of palisade and spongy tissue. According to our literature review, the average *S*_mes_ value is 15.9 m^2^ m^−2^, ranging from 2.8 m^2^ m^−2^ of *Arabidopsis cgr2/3* mutant leaf (Weraduwage *et al*., 2016) to 41.6 m^2^ m^−2^ of sun *Camellia japonica* leaf (Terashima *et al*., 2006). To facilitate CO_2_ diffusion in the leaf, most of the surface area of mesophyll cells are exposed to intercellular airspaces (Nobel, 2009). Greater total length of mesophyll cells exposed to intercellular airspaces thus is prerequisite to obtain the greater values of *S*_mes_. Terashima *et al*. (2011) illustrated four strategies of improving mesophyll surface area, including (1) cell elongation, (2) cell elongation accompanied by cell division, (3) decrease in cell size, (4) armed cells of grass species having lobes. For lobed cells or armed cells, high surface area exposure of mesophyll cells and chloroplasts to intercellular spaces and enhanced mesophyll conductance was found in *Oryza* spp and *Viburnum* spp (Sage and Sage, 2009; Chatelet *et al*., 2013). Adachi *et al*. (2013) and He *et al*. (2017) showed two rice lines named BTK-a and BTK-b to have 20-50% higher photosynthetic rate than the parents because of highly developed lobes of mesophyll cells. Although there are slight differences in the strategies to improve mesophyll surface area, the key of the other three approaches is the trade-off between cell size and cell density because the mesophyll cell size is negatively correlated with cell number when the leaf thickness is constrained (Pyankov *et al*., 1999).

To understand Smes easily, here some geometrical idealization is adopted. Let us assume that all mesophyll cells in a cross-section are spherical; cells are tightly packed and all cell walls are exposed to the intercellular airspaces (Fig. 3). Thus, Smes is equal to the total circumference of the spherical cells divided by the width of the cross-section according to the 2D model. If there is only a single uniform sphere in cross-section, *S*_mes_ is *π*. If cell diameter is reduced to one-third, three layers with nine uniform spherical cells would produce a *S*_mes_ of 3*π*. Increasing cell density would significantly improve *S*mes when the leaf thickness remains unchanged. Indeed, mesophyll cell density cannot be increased indefinitely. Smaller cell volume would result in over-lapping chloroplasts in mesophyll cells because of the strong correlation between the number of chloroplasts in a cell and the mesophyll cell area (Pyke and Leech, 1991, 1992). Therefore, the optimum is the minimum volume of mesophyll cell in which all the chloroplasts are tightly attached to cell wall and avoid over-lapping of chloroplasts. We have not rigorously examined the published photomicrographs of leaf cross-section, however, the number of chloroplasts in individual mesophyll cells seen in 2D cross-section is usually about 10-30. Compared with other leaf anatomical features, the variation in length and thickness of chloroplasts is smaller, with an average of 4.81 μm and 2.46 μm, respectively. The typical spherical mesophyll cell area containing 10-30 chloroplasts occupying all mesophyll surface area facing intercellular airspace is between 241-1780 μm^2^. If the mesophyll surface area is less than that value, chloroplasts might overlap in the mesophyll cell, which would result in the decrease of Sc. In addition, adequate mesophyll cell space is a prerequisite to achieve an increase in cell density. From Fig. 3, it could be found that fas is still 0.215, despite many more mesophyll cells. From the literature, the average *f*_ias_ is 0.241 regardless of species and environment, varying from 0.059 of *Oryza sativa* cslf6-2 mutant (Ellsworth *et al*., 2018) to 0.675 of *Asplenium scolopendrium* (Carriquí *et al*., 2015). Great variations in *f*_ias_ produce the possibility to utilize internal volume of the leaf to assemble more mesophyll cells to improve *S*_mes_ and *S*_c_. Authentic mesophyll cells are not uniformly spherical and are more complex, nevertheless, for most species the intercellular space is sufficient to be equipped with more mesophyll cells. Appropriate manipulation of mesophyll cell density to increase mesophyll and chloroplast surface area exposed to intercellular air space is a potential approach for enhancing leaf photosynthetic capacity by altering leaf anatomy.

**Fig. 3.**
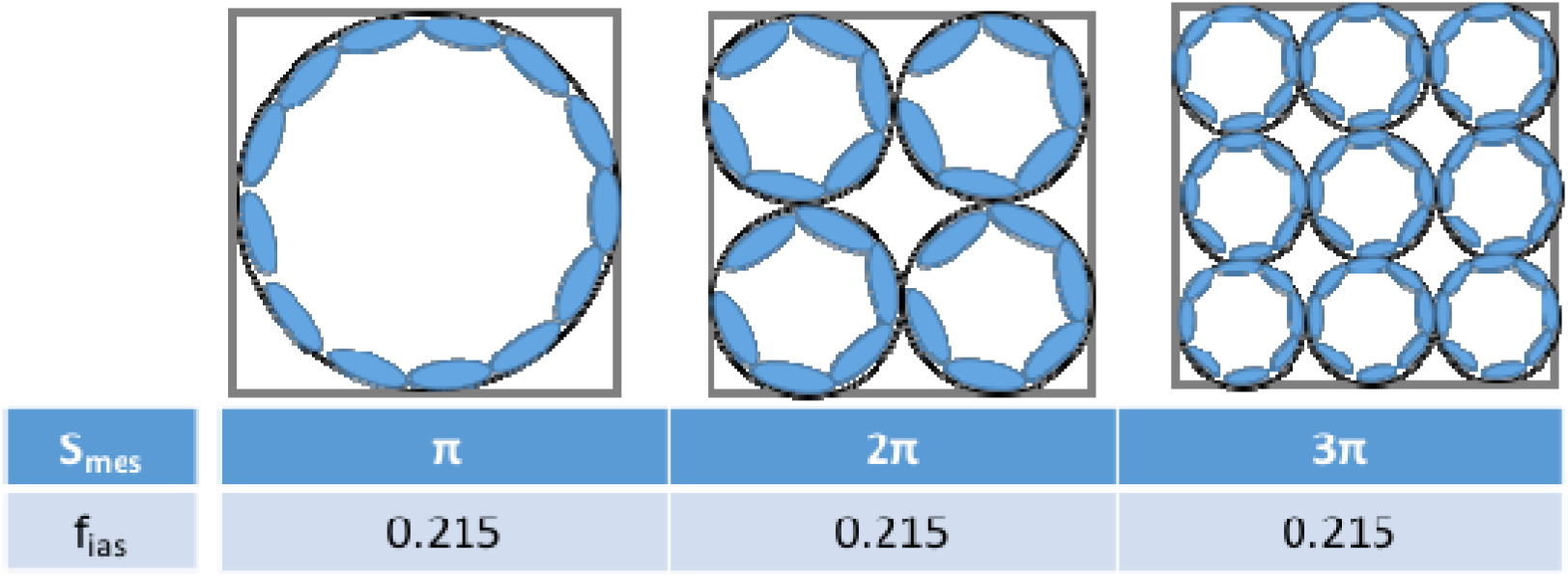
Representations of geometrical idealization of mesophyll cells showing how geometry affects the S_mes_ and f_ias_ value. *Smes*, mesophyll surface area exposed to intercellular air space; *fias*, fraction of intercellular air space.

## 3. Potential changes of photosynthetic capacity by manipulating mesophyll cell morphology

### 3.1 The mechanism of enhancing photosynthesis CO_2_ diffusion

After arriving at substomatal cavities, atmospheric CO_2_ diffuses through intercellular airspaces and then passes through the cell wall, cytosol, chloroplast envelope, finally reaching the sites of carboxylation in the chloroplast. The total CO_2_ diffusion resistance between substomotal cavities and carboxylation sites is termed mesophyll resistance and its reciprocal is defined as mesophyll conductance (*g_m_*), which is sufficiently small to restrict photosynthesis (Warren, 2008; Flexas *et al*., 2008, 2012; Evans *et al*., 2009). Mesophyll resistance can be divided into the gas-phase and liquid-phase resistance, in which liquid-phase resistance could account more than 90% of mesophyll resistance (Parkhurst, 1994; Tosens *et al*., 2012; Lu *et al*., 2016). In the liquid-phase, many CO_2_ diffusion parameters have not been accurately quantified (Evans *et al*., 2009; Xiao and Zhu, 2017), however, cell wall thickness and Sc are considered the most important anatomical parameters influencing gm (Syvertsen *et al*., 1995; Tomás *et al*., 2013). Summarized data confirmed the significantly negative correlation between *g*_m_ and cell wall thickness (Fig. 2). By contrast, *g*_m_ is positively correlated with *S*_c_ because a large chloroplast surface area adjacent to the cell wall is beneficial for fast diffusion through the cytosol and minimizes the total diffusion resistance. He *et al*. (2017) reported an increase in the number of lobed mesophyll cells induced much higher total surface area of mesophyll cells in the BTK lines, further resulting in greater mesophyll conductance. As shown in Fig.3, without increasing leaf thickness, increasing mesophyll cell number improves the mesophyll surface area, thereby increasing the number of chloroplasts per unit leaf area and *S*_c_. Higher *S*_c_ improves CO_2_ mesophyll conductance, increases CO_2_ concentration at carboxylation site and enhances photosynthetic capacity.

### Light interception and distribution in leaves

Leaf anatomy is well designed to capture solar and maximize light absorption; leaves often absorb more than 85% of the photosynthetically active radiation that hits them (Evans and Poorter, 2001). Lens-shaped epidermal cells can concentrate incident light severalfold, columnar palisade cells facilitate the penetration of light into the leaf, and randomly arranged spherical spongy cells and intercellular airspaces intensely scatter and reflect light, which increases the probability of light absorption for photosynthesis (Vogelmann *et al*., 1996a,b; Ustin *et al*., 2001; Smith *et al*., 2004). With depth into a leaf, the light and chlorophyll fluorescence intensity, especially for red and blue light, decline remarkably due to absorption and scattering, depending upon pigmentation as well as the mesophyll structure (Vogelmann, 1993; Vogelmann and Evans, 2002).

Cui *et al*. (1991) showed more and elongate palisade cells within sun leaves facilitated the light transmission in leaves and spread the light through the depth of the leaf while less compact palisade cells within shade leaves helps increase light absorption through greater scattering. Thus, altering leaf anatomy through regulating mesophyll cell geometry and packing would greatly influence the distribution of internal light. More and slightly-stacked mesophyll cell layers could efficiently facilitate the penetration of light into the leaf reducing light absorption in the top most layer that might be saturated by light allowing light to penetrate to the lower layers that may not be saturated by light. On the other hand, smaller intercellular airspace with small pockets could scatter light more (Smith *et al*., 2004), increase the light pathlength and enhance light absorption. Besides reducing CO_2_ diffusion resistance, higher *Sc* accompanied by more chloroplasts exposed to intercellular airspace aids efficient light absorbtion and helps satisfy the electron requirement for CO_2_ assimilation associated with high CO_2_ conductance (Terashima *et al*., 2009). The transgenic line *ATML1_pro_: KRP1* leaves exhibited an increase in palisade cell density improving air channel circularity and density, promoting the proportion of light energy used and operational election transport rate (Lehmeier *et al*., 2017).

### 3.2 Other potential changes

Leaf growth is determined by the amount of photosynthetic C partitioned between leaf area growth and *LMA* (Weraduwage *et al*., 2015). An alteration in C investment to leaf area or *LMA* could influence plant growth. Weraduwage *et al*. (2016) showed that overexpression of the *Cotton Golgi-related (CGR)* genes *CGR2* and *CGR3*, which mediate pectin methylesterification, increased C partitioned to leaf area growth and improved whole-plant photosynthesis and plant growth. Greater numbers of mesophyll cells would result in higher *LMA* (Pyankov *et al*., 1999; Poorter *et al*., 2009; John *et al*., 2017), which requires more C invested in *LMA* and reduces the available C for leaf area growth. Additionally, mesophyll cells make more than 90% of total leaf respiration owing to more mitochondria in mesophyll cells (Long *et al*., 2015a). More mesophyll cells mean more mitochondria and higher leaf respiration, reducing photosynthetic C available for plant growth (Weraduwage *et al*., 2016). Manipulating mesophyll cell density could be used to enhance photosynthetic capacity, nonetheless, it might change the allocation of photosynthetic C, further affecting plant growth.

Previous studies have described strong correlations between leaf hydraulic conductance and CO_2_ diffusion and photosynthesis rate (Flexas *et al*., 2013; Scoffoni *et al*., 2016; Barbour, 2017). Both leaf water transport and CO_2_ delivery have similarities in leaf anatomical traits. The leaf hydraulic system is composed of the major and minor veins, bundle sheaths, and the mesophyll cells outside the vein xylem. Mesophyll cell characteristics, such as mesophyll thickness, mesophyll cell porosity and connectivity, mesophyll cell surface area for evaporation, which significantly affect CO_2_ diffusion in leaf, are key parameters determining hydraulic conductance (Sack *et al*., 2015). Greater porosity of mesophyll cells contributes more to a high leaf hydraulic conductance, and broken connections between cells and fewer pathways of water flow would cause a lower aquaporin activity (Scoffoni *et al*., 2014). Small leaves with smaller cells tend to have greater minor vein length per unit leaf area, conferring high leaf hydraulic conductance (Scoffoni *et al*., 2011). Changes in gas exchange traits associated with altering mesophyll cell morphology would influence leaf hydraulic conductance and transpiration. Theoretically, increasing mesophyll cell density would not affect intercellular airspace (Fig. 3), nevertheless, an increase in mesophyll cell density inescapably decreases intercellular air space (Lehmeier *et al*., 2017). Hanba *et al*. (2004) documented that the leaves of transgenic rice Tr6322-H had higher mesophyll cell density and more developed collenchyma coupled to a higher mesophyll conductance, however, its leaf development might be influenced by water deficit because of increased water loss. The regulation of mesophyll cell morphology has potential to enhance photosynthetic capacity, however, it is worth studying whether it will increase hydraulic conductance simultaneously or change the composition of hydraulic conductivity.

Terashima *et al*. (2006) have discussed the importance of having sufficient *S*_c_ by decreasing cell size for efficient photosynthesis and compared the merits and demerits of different mesophyll cell sizes. Different species can have considerably distinct mesophyll cell sizes as a result of natural selection in various environments. Annual herbaceous plants often have large mesophyll cells, while ever-green trees tend to have smaller mesophyll cells. The leaves with small mesophyll cells have strong mechanical strength, long longevity, small heat capacity, and lower expansion rate. Additionally, they exhibit higher information cost (nuclear N- and P and chloroplast N and P). From this perspective, plants in natural environments may not benefit from very small mesophyll cells.

However, for agroecosystems, there might be more advantages to select small mesophyll cells. High mechanical strength would be beneficial to enhance crop pest resistance. The maintenance of leaf morphology depends on the cell wall, which is helpful to increase resistance to drought stress. For intensive agricultural production, N and P inputs are always sufficient or even excessive. If manipulating mesophyll cell morphology could improve photosynthetic capacity and increase crop yield without additional inputs, then N and P utilization efficiency would be higher. Selecting for smaller mesophyll cells does not mean completely changing the shape and size of the mesophyll cells. Instead, depending on the current size and arrangement of mesophyll cells, it may be possible to reduce mesophyll cell size appropriately by altering cell proliferation and expansion to improve *S*_c_ and reduce CO_2_ diffusion resistance, thereby enhancing photosynthetic efficiency. Indeed, comprehensive studies estimating the potential for regulating mesophyll cell morphology and the related changes on physiological and biochemical should be made.

## 4. Target genes could be used to regulate mesophyll cell morphology

Leaf growth has been divided into three phases: (1) leaf primordia occur at the shoot apical meristem, (2) cells proliferate and form specific structures during primary morphogenesis, (3) leaf cells begin to expand at secondary morphogenesis (Donnelly *et al*., 1999; Andriankaja *et al*., 2012). In the transition from primary to secondary morphogenesis, a retrograde chloroplast signal induces the onset of cell expansion and the leaf becomes photosyntheticaly active at the same time (Andriankaja *et al*., 2012), indicating that leaf growth as well as leaf morphology is inseparably correlated with photosynthesis. Although leaf morphology is regulated in response to different growth stages and environments, genetic control is fundamental to produce various leaves in nature. Thus, leaf morphogenesis and the related genes are hotspots in plant biology research. Numerous Arabidopsis mutants with altered leaf size and/or shape have been isolated (Horiguchi *et al*., 2006) and a key aspect of genes affecting leaf morphogenesis also have been identified during the past years. Several articles have reviewed in detail the critical genes regulating leaf shape, size, cell proliferation and expansion (Beemster *et al*., 2003; Micol, 2009; Horiguchi and Tsukaya, 2011; Gonzalez *et al*., 2012; Rodriguez *et al*., 2014, 2016; Hepworth and Lenhard, 2014). However, leaf development is a complex biological process and is determined by many genes and pathways. The molecular mechanisms underlying leaf size and shape are only starting to be unraveled. Although extensive research is still needed, key genes can provide some mechanisms for regulating leaf morphology.

Cell number, which plays a more important role in determining the final size of plant organs rather than cell size (Gázquez and Beemster, 2017), mainly depends on cell proliferation. Based on the existing research results, we divide the principal genetic factors regulating cell proliferation into three groups:

1. **Direct regulators.** Overexpression or loss of these genes can directly regulate cell proliferation. such as GROWTH-REGULATING FACTORs (*GRF*) family, which has been widely studied (Omidbakhshfard *et al*., 2015; Tsukaya, 2016). There are nine members in the *Arabidopsis thaliana* GRF family (AtGRF), of which AtGRF1, 2, 3, 4, 5, and 9 are involved in the regulation of cell proliferation. However, while other family members have partly overlapping roles, the function of *AtGRF5* is to regulate cell proliferation. When *AtGRF5* and *AN3* (the transcription coactivator) were overexpressed, plants developed larger leaves as a result of increasing cell number rather than cell size (Horiguchi *et al*., 2005). For AtGRFs, expression is regulated by miRNA miR396 and ectopic expression of miR396 repressed the AtGRF activity and resulted in smaller leaves with reduced cell number (Rodriguez *et al*., 2010). Additionally, overexpressing other genes, such as *AINTEGUMENTA (ANT)* (Mizukami and Fischer, 2000), *Arabidopsis SKP1-LIKE1* (ASK1) (Zhao *et al*., 1999), *Arabidopsis thaliana NGATHA* (AtNGA) (Lee *et al*., 2015), *G-PATCH DOMAIN PROTEIN1 (gdpl)* (Kojima *et al*., 2018), *KLU* (Anastasiou *et al*., 2007), *OLIGOCELLULA1, 4, 6 (olil, 4, 6)* (Fujikura *et al*., 2009), *POINTED FIRST LEAD 2* (*PFL2*) (Ito *et al*., 2000), *ROTUNDIFOLIA4* (*ROT4*) (Narita *et al*., 2004), *STRUWWELPETER* (SWP) (Autran *et al*., 2002) and *TCP4* (Schommer *et al*., 2014) could prolong cell proliferation and produce large leaves with more cells.
2. **Genes related to hormone signaling pathways.** Numerous hormones, such as auxins, brassinosteroids, cytokinin, gibberellins, jasmonic acid, etc, could act as signals for cell proliferation and expansion at the cellular level. The genes regulating hormone signals influence cell proliferation and expansion. For example, ectopic expression of *ARGOS* transduces auxin signals downstream of *AXR1*, which prolongs *ANT* expression and regulates cell proliferation, displaying enlarged leaves attributed to increase in cell number (Hu *et al*., 2003). The *cro* mutants *det2* and *dwf1*, affected in brassinosteroid biosynthesis revealed fewer and smaller cells (Nakaya *et al*., 2002). AHK-cytokinin receptor in *Arabidopsis thaliana* (Nishimura *et al*., 2004), PHYB-involved in gibberellin biosynthesis (Chaiwanon *et al*., 2016), and *aos (allene oxide synthase)* and *coil-16B* (*coronatine insensitive* 1)-jasmonate receptor, could influence cell division and expansion by regulating related hormone signals.
3. Genes related to pathways synthesizing structural component. Plant cells are encircled by a rigid cell wall and the biosynthesis and modification of cell wall would significantly influence cell expansion and proliferation. Cellulose, hemicellulose, and pectins are the dominant components of the plant cell (Ochoa-Villarreal *et al*., 2012; Tenhaken, 2015) and the genes code for enzymes catalyzing cellulose, hemicellulose, and pectin synthesis could regulate cell expansion and proliferation. For example, overexpression of the *fras5*, a mutant allele of *AtCesA7* gene involved in catalyzing cellulose synthesis, reduced cellulose content, cell wall thickness, and cell elongation (Zhong *et al*., 2003). The *ccr1* mutant has high ferulic acid content, an intermediate in lignin biosynthesis, and increased cell proliferation (Xue *et al*., 2015). A knockout of *CGR2* and *CGR3* genes showed reduced cell expansion and overall plant growth associated with reducing pectin methylesterification (Kim *et al*., 2015).

The directly regulated genes, which might be involved in signaling, structural, and other aspects regulating cell and leaf shape and still require further research, mainly regulate cell proliferation. In most instances, overexpressing these genes results in larger leaves owing to increased cell number rather than cell size. However, genes related to hormone signaling and structural component synthesis always influence both cell proliferation and cell expansion. Generally, an increase in the number of cells leads to a reduction in the volume of the cells, which is called “compensatory effect”. Horiguchi and Tsukaya (2011) defined compensation as when “a decrease in cell number in a leaf caused by genetic defect leads to an enhanced cell expansion”. They thought altered cell proliferation triggered compensation rather than cell expansion. Moreover, compensation is induced only when cell proliferation is beyond a threshold level (Horiguchi *et al*., 2006). It indicates increasing cell number within a certain range would not cause the changes in cell size, which provides the theoretical support of regulating mesophyll cell density to improve the *S*_mes_ and *S*_c_ values.

Besides manipulating mesophyll cell density, improving chloroplast density is helpful to increase the *S*_c_ values. Thus, the genes regulating chloroplast division will also be reviewed. The *ARC* (ACCUMULATION and REPLICATION of CHLOROPLASTS) mutants are an important genetic resource in the control of chloroplast division (Pyke and Leech, 1994; Gao *et al*., 2003; Cho *et al*., 2012; Zhou *et al*., 2015). Gao *et al*. (2003) revealed that mutatants of *ARC5* exhibit defects in chloroplast constriction and have enlarged dumbbell shaped chloroplasts. *GC1* (GIANT CHLOROPLAST) is another gene involved chloroplast division, which plays a positive role at an early stage of the division process, and *GC1* deficiency resulted in mesophyll cells harboring one to two giant chloroplasts (Maple *et al*., 2004). Hymus *et al*. (2013) showed overexpression of *ATHB17* increases chloroplast numbers per unit cell size due to an increase in the number of proplastids per meristematic cell. As one of the critical genes regulating cell proliferation, *AtGRF5* stimulates chloroplast division, resulting in a high chloroplast number per cell and a higher photosynthesis rate (Vercruyssen *et al*., 2015). Additionally, cytokinin promotes chloroplast ultrastructure and chlorophyll synthesis. Cortleven and Schmulling (2015) summarized cytokinin signaling pathways of Arabidopsis regulating chloroplast development and function through transcription factors. As the cytokinin receptors, *AHK2* and *AHK3* genes not only trigger cell divisions (Nishimura *et al*., 2004), but also regulate chloroplast development and chloroplast division. Recently, Larkin *et al*. (2016) described how the *REC* gene helps to establish the proportion of cellular volume devoted to chloroplasts; the *rec1* mutant has reduced size of the chloroplast compartment. Altered chloroplast coverage in cells might induce changes in *S_c_* and *g*_m_.

## 5. Prospects to manipulate mesophyll cell morphology for enhancing photosynthesis

Regulation is disconnected from limitation (Sharkey and Weise, 2012). Light and CO_2_ are the primary external constraints (limitations) on photosynthesis, therefore, regulating leaf morphology to take maximum advantage of light and CO_2_ is key to maximize photosynthesis. During the past decades, considerable research has been conducted to study the influence of leaf morphology on photosynthetic capacity in different species and environments. These studies help interpret how plants achieve high photosynthetic efficiency by optimizing leaf anatomy under different environments. Nonetheless, for any given species, the leaf architectural traits with the highest photosynthetic capacity is not always clear. Small differences in leaf phenotypes and the large number of genes that affect leaf architecture make research in this area difficult. As the most widely used species in plant biology research, *Arabidopsis thaliana* has been screened and numerous mutants with different leaf size and/or shape have been isolated (Horiguchi *et al*., 2006; Micol, 2009). A large number of the mutated genes determining cell proliferation, expansion and differentiation have been identified. However, these mutations have not typically been studied in the context of the effects on photosynthesis. Therefore, the best targets for engineering leaf anatomy are unknown. More, and in-depth, studies should focus on (1) the differences in photosynthetic capacity under different leaf shape and size as well as leaf anatomy, (2) how to influence leaf photosynthesis through the potential target genes altering leaf morphology. Such research will pinpoint the targets for altering leaf architecture to maximize photosynthetic capacity.

Although light fluctuations significantly influence leaf photosynthesis (Kaiser *et al*., 2018; Slattery *et al*., 2018), CO_2_ concentration is the dominant factor restricting photosynthesis and yield in crops under current conditions (Ainsworth and Long, 2005; Ellsworth *et al*., 2012). In C_4_ plants, the specialized Kranz anatomy helps concentrate CO_2_ concentration and enhance photosynthetic efficiency. C_3_ plants do not possess Kranz anatomy and CO_2_ diffusion resistance in mesophyll cells significantly reduces the CO_2_ concentration at the sites of carboxylation. If mesophyll resistance could be eliminated, photosynthesis rates would be improved by up to 20% (Zhu *et al*., 2010). Increasing chloroplast area exposed to intercellular airspace as much as possible could reduce the CO_2_ diffusion resistance and enhance mesophyll conductance and photosynthetic capacity. Terashima *et al*. (2001, 2006, 2011) have elaborated the importance of having sufficient *S*_c_ for efficient photosynthesis and have discussed the various strategies to increase the *S*_mes_ and *S*_c_ values by manipulating cell proliferation and expansion. They thought decreasing mesophyll cell size was an effective approach to increase *S*_c_ and decrease CO_2_ diffusion resistance. Nevertheless, most plants do not tend to form small mesophyll cells as a result of natural selection. As early as the 1970s, selection for a smaller cell size have been considered as critical parameters to breed new variety with high photosynthesis and yield. Nevertheless, owing to poor understanding of molecular mechanisms controlling leaf anatomy as well as cell proliferation and expansion, the investigations into the influence of mesophyll cell size on photosynthesis have not been fully developed and the selection of more and smaller mesophyll cells is not currently considered a potential approach to enhance photosynthesis capacity. Recent research shows increasing mesophyll cell number through regulating genes related to the cell cycle are beneficial for improving photosynthetic capacity (Takai *et al*., 2013; Lehmeier *et al*., 2017). It can be foreseen that engineering leaf anatomy by manipulating cell proliferation and expansion using gene regulation or gene editing technology will be a potential approach to enhance plant photosynthetic capacity in future.

Indeed, many challenges, including photosynthesis and molecular mechanisms controlling leaf anatomy, need to be solved. What is the potential for regulating cell density to maximize photosynthesis? As mentioned in the compensation effect, increasing in mesophyll cell number beyond a threshold value will trigger a decrease in cell size (Horiguchi *et al*., 2006). Smaller mesophyll cell size will restrict the number and division of chloroplasts. Although the *S*mes value is improved by increasing mesophyll cell number, the Sc value might not be increased because of over-lapping chloroplasts, which is unfavorable for enhancing photosynthesis. It is therefore essential to explore the relationship between mesophyll cell traits and mesophyll conductance and photosynthetic capacity. Additionally, an increase in leaf mesophyll cell density would improve leaf mass density and *LMA*, which could modify photosynthetic C partitioning to leaf area growth and growth in terms of *LMA*, further influencing plant growth. Weraduwage *et al*. (2016) revealed that in the *cgr2/3* mutant greater C partitioning to *LMA*, associated with an increase in leaf cell density, resulted in smaller leaf area and reduced whole-plant photosynthesis. A great number of cells and organelles require more energy and hence incur higher maintenance respiratory costs (Weraduwage *et al*., 2016). What’s more, alteration in mesophyll cell number and size also would impact the interception of light, water transport and CO_2_ assimilation. Comprehensive studies to analyze the effects of manipulating mesophyll cell morphology on other physiological processes in leaves would be useful to optimize the approach for enhancing photosynthetic capacity.

The selection of target genes for altering leaf anatomy also requires more accurate and extensive research. As mentioned in Lehmeier *et al*. (2017), the suppression of *RBR1* removes some restrictions on cell division, leading to more and smaller mesophyll cells; however, the transgenic lines with higher photosynthetic capacity grew smaller than the wild type because of higher *LMA*. Although photosynthetic capacity is improved by regulating these genes, such plants do not achieve higher plant biomass and crop yield as result of the decrease of photosynthetic area. The genes coordinating the improvement of photosynthetic capacity and plant growth can meet the demand of modern agriculture for high yield.

Altering leaf anatomy to enhance leaf photosynthetic capacity must be achieved without influencing leaf area. From this perspective, the genes directly regulating cell proliferation might be considered first. For example, overexpression *AtGRF5* and *AN3* in *Arabidopsis thaliana* results in larger leaves with an increased cell number (Horiguchi *et al*., 2005). Additionally, the genes controlling cell proliferation and expansion might change other physiological processes. For example, the *cgr2/3* mutant, double knockout of *CGR2* and *CGR3* genes, showed a reduced cell expansion and increased mesophyll cell density as result of a reduction in the degree of pectin methylation; however, it induced hardening of cell walls, and increased CO_2_ resistance of the cell wall and reduced photosynthetic efficiency (Weraduwage *et al*., 2016). Altering expression of ANT genes modified cell proliferation and regulated the number of higher order veins and minor tertiary veins (Kang *et al*., 2007), which are closely related to leaf photosynthetic capacity (Sack *et al*., 2013; Feldman *et al*., 2017). It is therefore essential to understand the various functions of the target genes for accurate editing the genes to alter leaf morphology and enhance photosynthetic efficiency.

## 6. Conclusion

Natural selection allows for a range of leaf architectures to effectively capture light and exchange gases, resulting in high photosynthetic efficiency to grow and survive. Despite the varitety of species and environmental conditions, Sc is the most important indicator of leaf photosynthetic capacity. Having a sufficiently high *S*_c_ is a prerequisite of efficient photosynthesis. Chloroplast number and distribution in mesophyll cells and *S*_mes_ significantly influence *S*_c_. Without overlapping mesophyll cells, increasing mesophyll cell density is an important strategy to improve *S*_mes_ and *S*_c_. Depending on the manipulation of target genes involved in cell proliferation and expansion, such as *AtGRF* family, it should be possible to produce more mesophyll cells without influencing leaf area, further inducing higher *S*_c_ values and photosynthetic capacity. It is foreseen that the chain “gene regulation-leaf anatomy-photosynthetic capacity” will be an important strategy for enhancing photosynthetic capacity and crop yield. Indeed, there are many challenges that need to be thoroughly studied, such as the selection of target gene regulating cell proliferation and expansion, the potential of adjusting mesophyll cell number and size, and the effects of leaf anatomical traits on other physiological processes, including light utilization, water transport etc.

## Acknowledgements

This research was funded by U.S. Department of Energy Grant DE-FG02-91ER2002. Partial salary support for T.D.S. came from Michigan AgBioResearch. T.R acknowledges the fellowship supported by China Scholarship Council.

## List of Abbreviations

*A*_n_: , photosynthesis rate;
*f*_ias_: , fraction of intercellular airspace;
*g*_m_: , mesophyll conductance;
*g*_s_: , stomatal conductance;
*L*_chl_: , chloroplast length;
LMA: , leaf mass per unit leaf area;
*S*_c_: , chloroplast surface area exposed to intercellular airspace per unit leaf area;
*S*_mes_: , mesophyll surface area exposed to intercellular airspace;
*T*_chl_: , chloroplast thickness
*T*_cw_: , mesophyll cell wall thickness;
*T*_L_: , leaf thickness;
*T*_mes_: , mesophyll layer thickness.

